# Inferring the mode and strength of ongoing selection

**DOI:** 10.1101/2021.10.08.463705

**Authors:** Gustavo V. Barroso, Kirk E. Lohmueller

**Author notes:** **AUTHOR CONTRIBUTIONS:** GVB and KEL conceived and designed the study and wrote the manuscript. GVB conducted the simulations and implemented the method. **COMPETING INTERESTS:** The authors declare no competing financial interests.

## Abstract

Genome sequence data is no longer scarce. The UK Biobank alone comprises 200,000 individual genomes, with more on the way, leading the field of human genetics towards sequencing entire populations. Within the next decades, other model organisms will follow suit, especially domesticated species such as crops and livestock. Having sequences from most individuals in a population will present new challenges for using these data to improve health and agriculture in the pursuit of a sustainable future. Existing population genetic methods are designed to model hundreds of randomly sampled sequences, but are not optimized for extracting the information contained in the larger and richer datasets that are beginning to emerge, with thousands of closely related individuals. Here we develop a new method called TIDES (**T**rio-based **I**nference of **D**ominanc**e** and **S**election) that uses data from tens of thousands of family trios to make inferences about natural selection acting in a single generation. TIDES further improves on the state-of-the-art by making no assumptions regarding demography, linkage or dominance. We discuss how our method paves the way for studying natural selection from new angles.

## INTRODUCTION

Genetic variation in natural populations is balanced among mutation, natural selection, and genetic drift. Whether most mutations in genomes are deleterious, neutral, or beneficial has been debated throughout the history of population genetics, and remains controversial (Hey 1999; Kern and Hahn 2018; Jensen et al. 2019). One challenge in addressing this topic is that genetic diversity from natural populations likely has been affected by multiple evolutionary forces simultaneously, making it hard to isolate and test for the effects of any one such force (Barroso and Dutheil 2021). However, many non-synonymous mutations are detrimental (Keightley and Lynch 2003) and since natural selection has limited efficiency in removing them from the population, segregation of deleterious polymorphism is unavoidable, with direct consequences on the fitness of individuals (Agrawal and Whitlock 2011). In humans, for example, this genetic load contributes to a ∼50% miscarriage rate (Rice 2018), most of which occur in the first 12 weeks of pregnancy. Inferring the strength of selection is therefore a central goal in biology (Nielsen et al. 2007; Eyre-Walker and Keightley 2007). Equally important is characterizing the degree to which deleterious variants influence fitness when in heterozygous state, referred to as the dominance effect (Agrawal and Whitlock 2011; Huber et al. 2018). Together, knowledge of the selection coefficient (*s*) and the dominance coefficient (*h*) can inform about functional constraints in protein structure (Moutinho, Trancoso, and Dutheil 2019) and interaction networks (Park, Hescott, and Slonim 2019; Ratnakumar et al. 2020), shedding light into both evolutionary and medical genetics.

Existing methods to infer the strength of selection use the site frequency spectrum (SFS) to infer a distribution of fitness effects (DFE) of new mutations (Eyre-Walker, Woolfit, and Phelps 2006; Boyko et al. 2008; Kim, Huber, and Lohmueller 2017; Tataru et al. 2017; Keightley and Eyre-Walker 2007). Although they have greatly contributed to advancing population genetics in the past 15 years (Moutinho, Bataillon, and Dutheil 2020), these models have important shortcomings, stemming mostly from the limited amount of information retained in the SFS. First, they neglect linkage disequilibrium (LD) and selective interference among sites (Hill and Robertson 1966; Garcia and Lohmueller 2020). Second, they only incorporate oversimplified demographic histories. Third, the SFS alone cannot distinguish between *s* and *h* (Huber et al. 2018) and consequently the DFE is typically inferred assuming additivity – which is problematic since deleterious mutations tend to be recessive (Huber et al. 2018; Bosse et al. 2018). Fourth – and perhaps most importantly – the magnitude of selection can change over time (Wittmann et al. 2017; Orr and Betancourt 2001). For example, genes that have been highly constrained in the past may experience relaxation upon environmental change or even become positively selected. Conversely, previously neutral alleles may become deleterious. Several methods model a DFE with a proportion of positively selected variants (Galtier 2016; Schneider et al. 2011; Boyko et al. 2008; Zhen et al. 2021) but these methods still capture long-term signals of selection from the SFS, effectively averaging *s* over several thousand generations. Such averaging may result in misleading inference if there have been substantial fluctuations in selective pressures during the history of the population under study. Although progress has been recently made in inferring temporal trajectories of selection using ancient DNA samples (Mathieson 2020), a sufficient quantity of ancient samples is not always available from the relevant population. Taken together, these limitations reduce their ability to accurately infer selection occurring in contemporary populations.

Here we suggest a way forward to overcome these challenges in inferring fitness effects for mutations that takes a different perspective of modeling natural selection acting on a single-generation, rather than throughout the entire history of the population. Our new approach is made possible by the explosion of genome re-sequencing data from natural populations. Largely due to the medical implications, humans have been the organism benefiting the most from this data deluge. For example, deCode genetics has genotype data on more than 160,000 individuals and 60,000 whole-genome sequences of much of the population of Iceland, including 2,926 family trios (Halldorsson et al. 2019). Other projects like the UK Biobank have genotype data on 500,000 individuals, 200,000 exome sequences and project 200,000 whole-genome sequences by the end of 2021, while the TopMed project has sequenced 53,835 (Taliun et al. 2021), including 1,465 family trios (Kessler et al. 2020). It is anticipated that within the next few years, entire populations will be sequenced, incorporating hundreds of thousands of parent-offspring trios.

The availability of sequences from thousands of closely related individuals presents both an opportunity and a challenge. One the one hand, these data will provide an opportunity to overcome the aforementioned limitations of SFS-based methods in the inference of selection. On the other hand, these data present methodological challenges, as traditional assumptions (*e*.*g*., sampled individuals are unrelated) break down with such large samples. Further, such datasets require dedicated models that are accurate and computationally efficient. Here we overcome this challenge and develop a new model called TIDES (**T**rio-based **I**nference of **D**ominanc**e** and **S**election) that is able to infer dominance and natural selection using tens of thousands of parent-offspring trios. Our method is designed to handle such large datasets efficiently, in anticipation of their availability in the near future. Moreover, a unique feature of TIDES is that it is sensitive to the strength of selection acting on the current generation, and it is therefore ideal to study population-specific signatures of selection, while not being confounded by other evolutionary forces like demography, or averaging selective effects over long time periods. TIDES can be applied to either sets of variants across the genome or a single variant at a time, further showcasing its flexibility.

## RESULTS

### Overview of the model

TIDES uses approximate Bayesian computation (ABC) (Pritchard et al. 1999; Beaumont, Zhang, and Balding 2002; Beaumont 2019) to model the effect of selection on genetic diversity during the span of a single generation. It leverages phased sequences from parent-offspring trios to detect signatures of selection in the transmission of single nucleotide polymorphisms (SNPs) (Meyer et al. 2012) and uses this information to infer dominance and selection coefficients. By exploiting the random shuffling of haplotypes during meiosis, family trio data becomes immune to external confounding factors that lurk in traditional population genetic studies (Bates et al. 2020), such as non-equilibrium demography (Sul, Martin, and Eskin 2018; Barton, Hermisson, and Nordborg 2019). The first step in our simulation framework is to use parental haplotypes and recombination maps to generate an array of potential zygotes for each trio (**Figure 1A, Algorithm 1**). We then sequentially impose rounds of viability selection on the simulated zygotes, for independent values of *s* and *h* drawn from their prior distributions, and compute summary statistics from the set of “selected” zygotes (**Figure 1B**). Comparing genomes from children (the observed data) with genomes from simulated zygotes that could have been conceived by their parents provides information about the (unobserved) embryos that did not survive, and is therefore indicative of the strength of selection.

**Figure 1:**
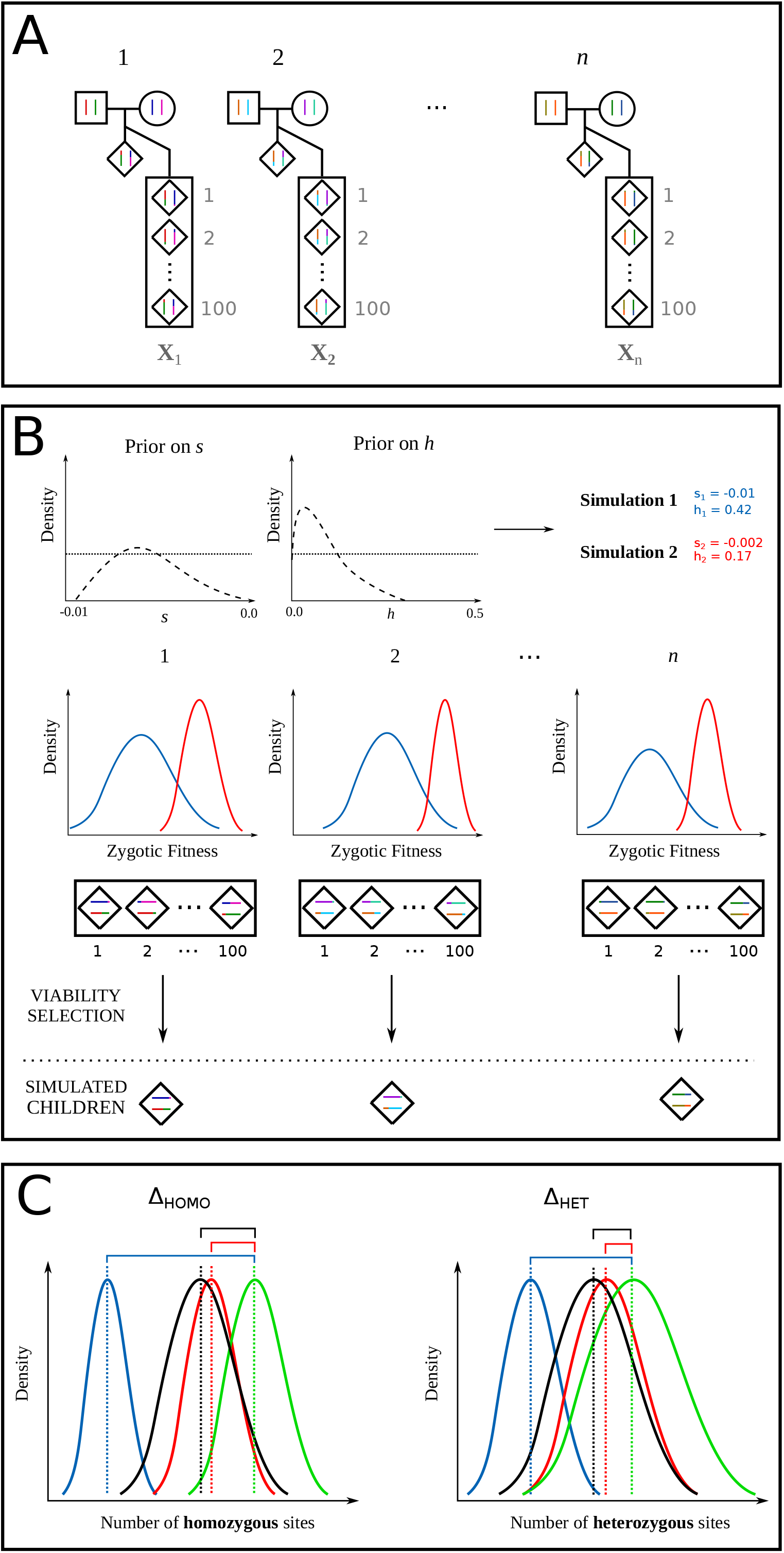
Schematic representation of TIDES. A: Observed family trios (black outline) together with the zygotes generated from parental haplotypes (gray outline). B: Illustration of the TIDES simulation engine for two draws from the prior distribution of *s* and *h* representing strong (blue) and weak (red) values of selection. The middle row shows the computation of zygotic fitness and natural selection. C: Comparison between the observed summary statistics (green) and the summary statistics from the simulations using the selection parameters from the prior distribution (red and blue). The left panel shows the comparison of the number of homozygous genotypes (Δ_HOMO_) and the right panel shows the comparison for the number of heterozygous genotypes (Δ_HET_).

In essence, TIDES is a fitness-based model that mimics the process of meiosis followed by natural selection. We compute the fitness *f* of each individual (both real and simulated) using a multiplicative model

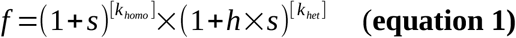

where **k**_**homo**_ and **k**_**het**_ are the counts of derived sites in homozygous and heterozygous states, respectively. To compute summary statistics, we consider the combined genetic diversity of the parents separately from the combined genetic diversity of the offspring. We use the relative differences in the averages of **k**_**homo**_ and **k**_**het**_ between offspring and parents as our two summary statistics, denoted **Δ**_**homo**_ and **Δ**_**het**_. Specifically, let **x**_**0**_, **y**_**0**_ be the average number of homozygous and heterozygous sites among parents, and **x**_**1**_, **y**_**1**_ be the corresponding averages among children. We define **Q**_**obs**_ as the vector storing the dimensionless quantities **Δ**_**homo**_ **=** (**x**_**1**_ – **x**_**0**_) / **x**_**0**_ and **Δ**_**het**_ = (**y**_**1**_ – **y**_**0**_) / **y**_**0**_. Likewise, if **x**^**(i)**^_**2**_ and **y**^**(i)**^_**2**_ are the averages among simulated zygotes that have survived selection for the *i*th parameter combination, then **Q**^**(i)**^_**sim**_ is the vector storing **Δ**_**homo**_ = (**x**^**(i)**^_**2**_ – **x**_**0**_) / **x**_**0**_ and **Δ**_**het**_ = (**y**^**(i)**^_**2**_ – **y**_**0**_) / **y**_**0**_. Disregarding *de novo* mutations, such trio-based summary statistics can be used to model both negative and positive selection on a set of candidate sites. When modeling negative selection, higher values of |*s*| (corresponding to stronger negative selection) should lead to sharper reductions in the overall number of deleterious variants in the offspring and therefore lower values of both **Δ**_**homo**_ and **Δ**_**het**_. Higher values of *h* should result in lower values of **Δ**_**het**_ but not **Δ**_**homo**_. Conversely, when modeling positive selection, higher values of *s* (corresponding to stronger positive selection) should lead to sharper increases in the overall number of beneficial variants in the offspring and therefore higher values of both **Δ**_**homo**_ and **Δ**_**het**_. Higher values of *h* should result in higher values of **Δ**_**het**_ but not **Δ**_**homo**_. Therefore, retaining genotype information instead of reducing genetic diversity to the SFS is key to disentangling the combined effects of dominance and selection.

### Evaluating the performance of TIDES

#### Inferring the exome-wide strength of negative selection

We first evaluate TIDES’ ability to infer the strength of negative selection on a set of deleterious SNPs. To benchmark our model and method, we simulated trio sequence data from a population reflecting the European demographic history (Gravel et al. 2011) and sex-specific recombination maps (Halldorsson et al. 2019), as well as the human exome structure using SLiM (Haller and Messer 2018) (Methods). These simulated data should reflect a reasonable picture of deleterious standing variation, both in terms of derived allele frequencies and their linkage disequilibrium patterns. We then sampled 60,000 individuals from the final generation, matched females and males at random to generate offspring and finally down-sampled to 50,000 trios in each scenario, which became our test data sets for inference. Throughout the following simulation study we employed flat, un-informative priors in order to assess TIDES’s ability to extract information from the data when there is weak *a priori* knowledge about the parameters. Specifically, we let *h* be uniformly distributed in the open interval (−0.1; 0.6) and *s* be log-uniformly distributed in the open interval (−10^−5^ ; -10^−1^) (except for neutral simulations, where *s* was uniformly distributed in the open interval (−0.01; 0.01)). The ranges of the priors reflect values of *s* and *h* that are pertinent based on the population genetics literature. The log-uniform prior on *s* was chosen such that the parameter space reflects the “magnitude” of selection, implying even exploration of values around -0.01 and -0.0001, for example.

We found that in neutral simulations, TIDES infers *s* to be tightly centered around zero (**Figure S1**), indicating that noise in the sampling of parental SNPs (*i*.*e*., genetic drift) does not generate a spurious signal of selection. TIDES has overall high accuracy in the six combinations of *s* (−10^−4^, -10^−3^, -10^−2^) and *h* (0, 0.5) that we tested, with the medians of the inferred posterior distributions centered around the true values of *s* (**Figure 2**). In general, precision is higher in the recessive scenarios and especially as selection becomes stronger. This change in power occurs because we treat our sample of parents as a *de facto* population, and project it forward by one generation where (disregarding *de novo* mutations) negative selection has the opportunity to reduce deleterious genetic diversity. Therefore, similarly to the classic population genetics result where selection becomes more efficient as |Ne * *s*| grows beyond 1 (Kimura 1979), in TIDES, the product of *n* (the number of trios) and *s* must be large enough to have high inferential power. Indeed, for our sample size of 50,000 trios, |*n* * *s*| equals 500 in the strong selection scenario and 50 in the moderate selection scenario, but only 5 when selection is weak. Taken together, these results suggest that for sufficiently large sample sizes, our trio-based framework can accurately infer the strength of ongoing selection for an arbitrary range of selection coefficients.

**Figure 2:**
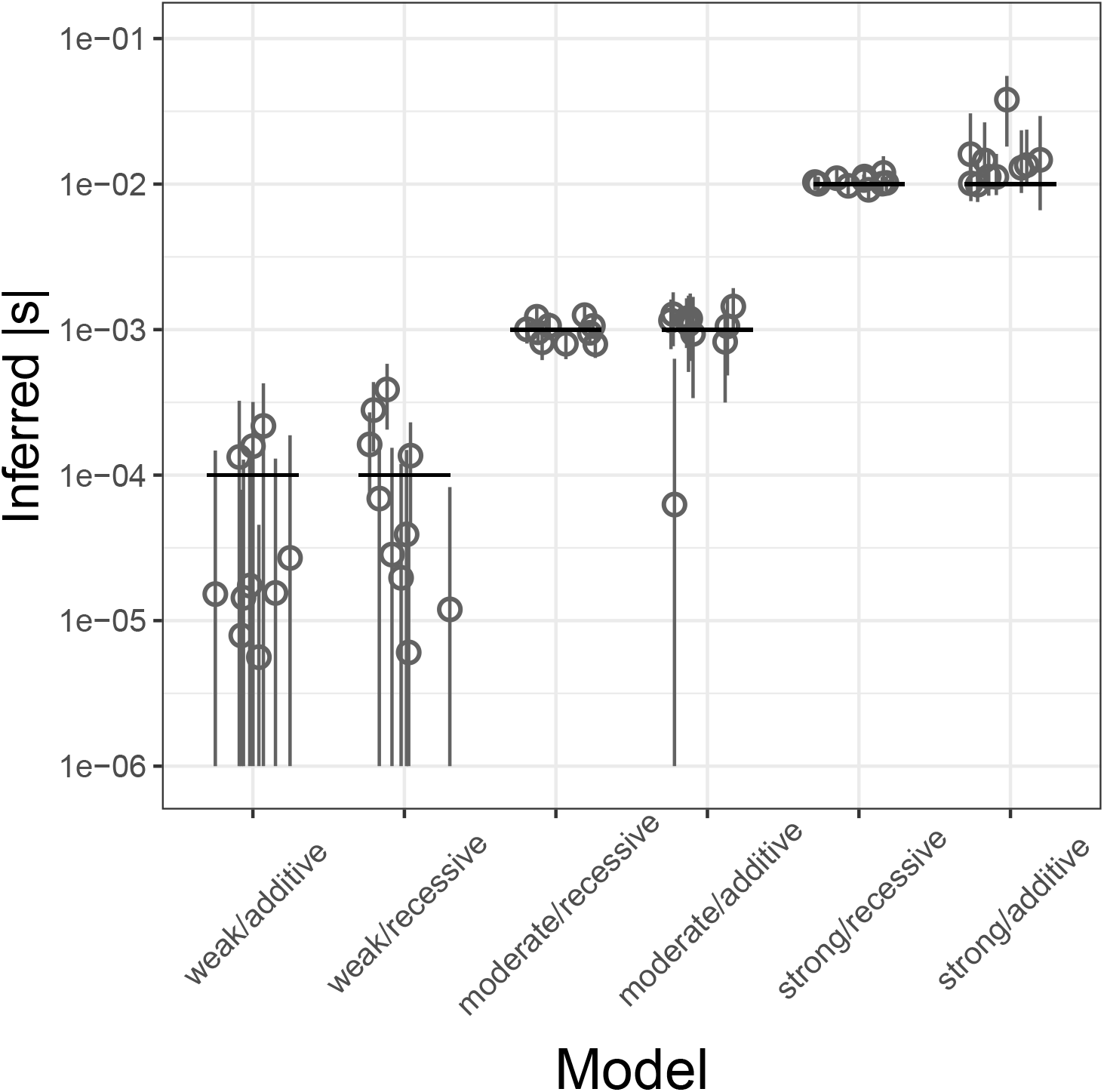
Inference of *s* from a genome-wide set of deleterious SNPs for different strengths of selection and dominance effects. Each scenario includes the estimates from 10 simulated datasets. True values are shown as black horizontal segments, with medians of the inferred posterior distributions denoted by gray circles and their 95% credible intervals by gray vertical lines. Y-axis in log_10_ scale, all values in absolute numbers. Here 50,000 trios are used.

It has been well established that the selection coefficients of deleterious SNPs vary by several orders of magnitude, from nearly neutral to lethal (Eyre-Walker, Woolfit, and Phelps 2006; Boyko et al. 2008; Kim, Huber, and Lohmueller 2017). To assess the performance of TIDES in the presence of a DFE, we performed simulations where the selection coefficient of new mutations comes from a Gamma-distributed DFE parameterized by α = -0.186 and β = 0.071. As negative selection purges strongly deleterious alleles more efficiently, the average selection coefficient of segregating SNPs is expected to be less negative than that of the DFE of new mutations. In our simulations of the European evolutionary history, the average *s* of new mutations is -0.013 whereas that of segregating SNPs is - 0.00015. Since TIDES focuses on a single generation instead of modeling long-term frequency trajectories, it infers the average *s* of SNPs segregating in the parents. Although the value of -0.00015 falls within the weak selection regime where our power with 50,000 trios is reduced, the medians of the posterior distributions are located near the true value (**Figure 3A**), showing that when mutations have different selection coefficients, TIDES can infer their average.

**Figure 3:**
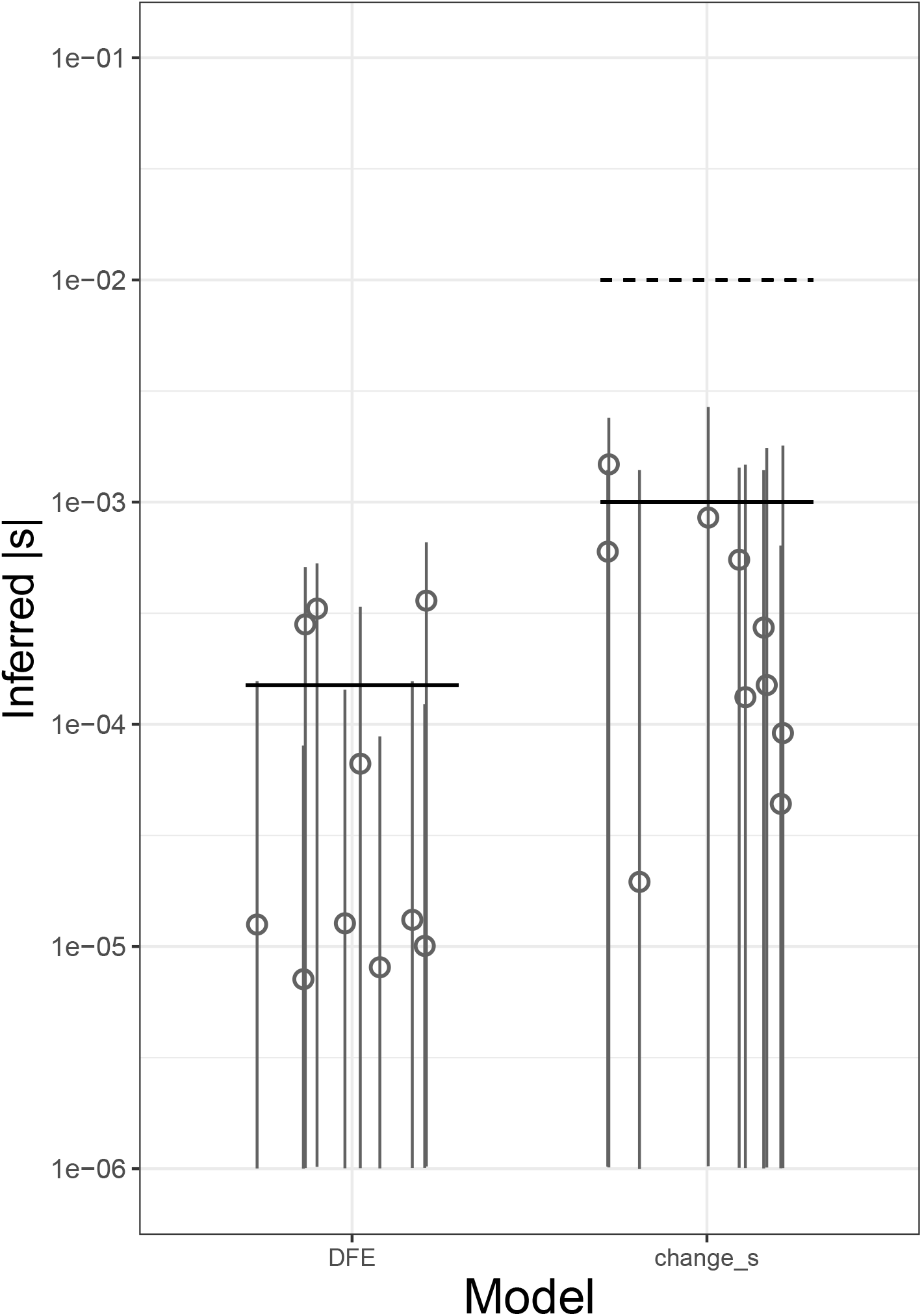
Inference *s* from a genome-wide set of deleterious SNPs under complex models of selection. Each scenario includes the estimates from 10 simulated datasets. Left panel shows the results when the true DFE follows a gamma distribution (mean value of *s* shown by black horizontal line). The right panel shows the case where there was a 10-fold reduction in the selection coefficient. Ancient and current values of *s* shown by dashed and solid horizontal lines, respectively. Medians of the inferred posterior distributions denoted by gray circles and their 95% credible intervals by gray vertical lines. Y-axis in log_10_ scale, all values in absolute numbers. Here 50,000 trios are used.

Finally, we asked whether TIDES could capture a shift in the strength of selection happening in current generation. The goal of this simulation was to mimic a scenario of environmental change where the selective pressure is abruptly reduced, and to assess the performance of our method in these data. To this end, we started from the same parental sequences from the recessive scenario with historical *s* = - 10^−2^ except this time we imposed viability selection in the offspring generation by changing *s* of all segregating SNPs to -10^−3^. Although a 10-fold decrease in the strength of selection is a statistically challenging signal to capture (because the level of standing variation reflects the previously stronger selection and is therefore reduced relative to the new expectation, with respect to both the number of SNPs and their frequencies), TIDES recovers the updated value of *s* (**Figure 3B**), showcasing that our model is sensitive to the strength of ongoing selection and is not burdened by memory of the past.

#### Inferring the selection coefficient of single deleterious SNPs

The results above suggest that TIDES can infer the (average) selection coefficient from a set of deleterious SNPs, but in some situations, single SNPs may be of interest. A few methods have been recently developed to infer selection coefficients of single SNPs using contemporary data (Stern, Wilton, and Nielsen 2019) or their temporal trajectories using ancient DNA data (Mathieson 2020), but tools for detecting ongoing selection are still lacking. To achieve high accuracy with single variants in TIDES, we once again should require that |*n* * *s*| *>>* 1, noting that only informative trios (those where at least one parent is heterozygous, hence the couple has the potential to produce more than one kind of offspring) should be included in the analysis. Because interest in individual SNPs may be motivated by situations where large effects are expected, we tested TIDES’s accuracy to infer strong negative selection using the open interval (−10^−4^ ; -10^0^) as a log-uniform prior on *s*. We simulated datasets where the frequency *q* of the deleterious allele among parents is ∼0.5, and we varied the number of informative trios (10,000, 30,000 or 100,000), as well as *s* (−0.01, -0.05 or -0.1) and *h* (0.0 or 0.5). TIDES is accurate in all scenarios of *s* = -0.1, whereas it requires at least 30,000 informative trios to have high accuracy when *s* = -0.05 and does not start to perform well until 100,000 trios and recessive selection for *s* = -0.01 (**Figure 4**). This means that unsurprisingly, TIDES’ accuracy on single SNPs depends on higher values of |*n* * *s*| than for exome-wide inference (since all couples are highly informative in the latter case due to the higher number of SNPs they carry), but that given enough data our model is able to infer ongoing selection on a single deleterious variant.

**Figure 4:**
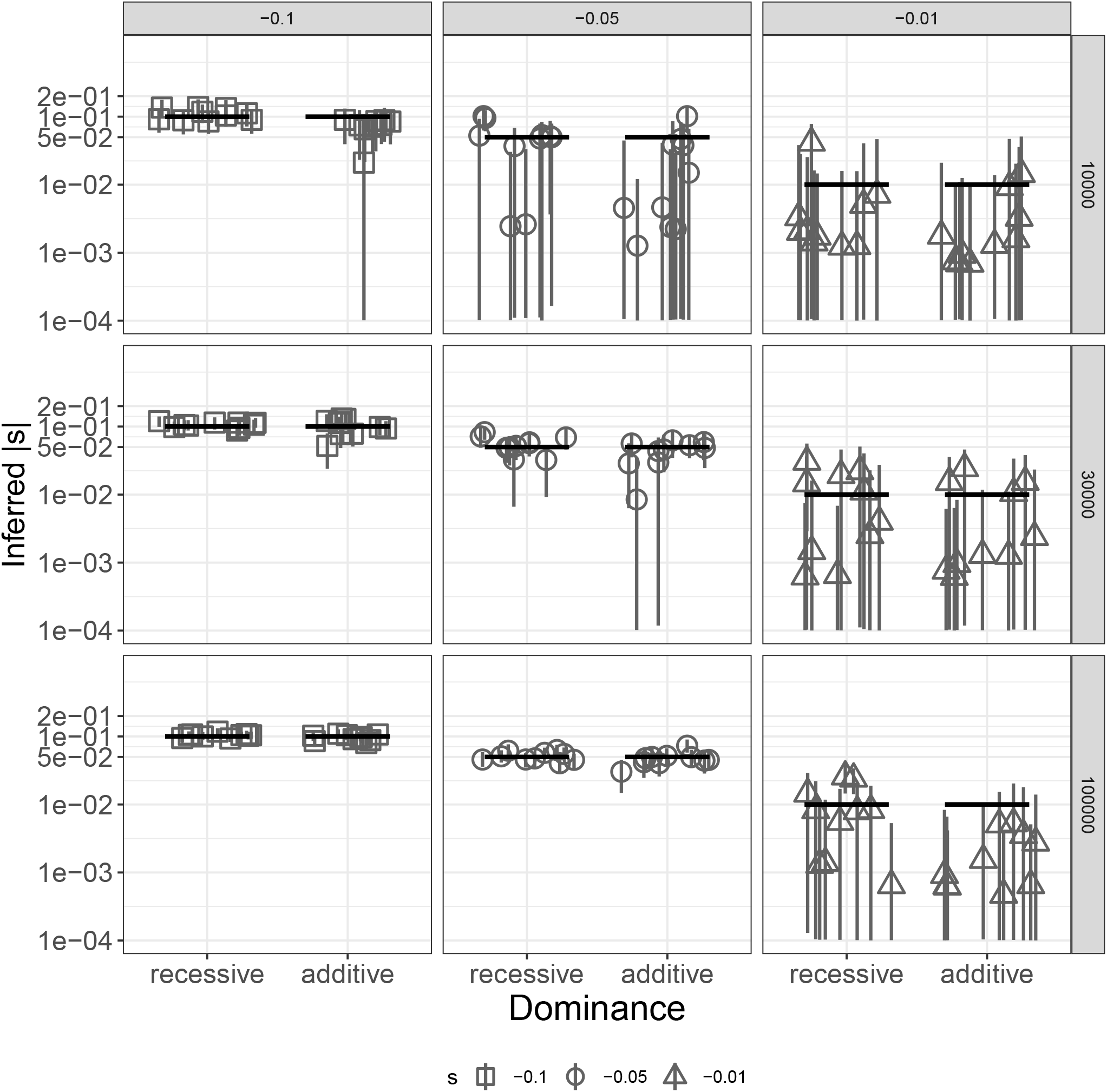
Inference of *s* for a single deleterious SNP with different dominance effects. Each scenario includes the estimates from 10 simulated datasets. Columns show different values of the true selection coefficient, and rows show different sample sizes, in terms of the number of trios used. True values are shown as black horizontal segments, with medians of the inferred posterior distributions denoted by gray shapes and their 95% credible intervals by gray vertical lines. Y-axis in log_10_ scale, all values in absolute numbers.

Since the frequency of deleterious SNPs segregating in natural populations is inversely proportional to the strength of negative selection against them, more strongly deleterious variants will tend to be kept at lower frequency, requiring larger samples from the population in order to find a sufficient number of informative trios. On the other hand, the number of informative trios required for inference decreases with increasing deleteriousness of variants because in this case each SNPs exerts a stronger signal in the data (*i*.*e*., sharper transmission distortion). Therefore, the overall sample size required for accurate inference (with subsequent down-sampling to consider only informative trios) is a function of both of these parameters that act on opposing directions. To assess whether it is realistic to expect that any single, strongly deleterious variant segregates at appreciable frequencies in humans, we performed simulations with exponential population growth where the final population size is 10,000,000, and assumed that the total number of target sites subject to mutations of each selection coefficient is 1,000,000 for with *s* = -0.1; 5,000,000 for *s* = -0.05; and 5,000,000 for *s* = -0.01 (Methods). In all cases, we found SNPs segregating at the absolute frequency thresholds required for accurate inference with TIDES (**Table 1**). In other words, this simulation shows that we would expect a few strongly deleterious SNPs to segregate at frequencies high enough such that it would be possible to subset a collection of 10,000,000 trios down to a sample size where both *q ∼* 0.5 and we meet the required number of informative trios. We conclude that there are strongly deleterious variants likely segregating in the population at sufficient frequency to be analyzed using TIDES in the foreseeable future. With the sample sizes described here, it will already be possible to use the single-variant model in TIDES to test for strong ongoing selection, which can provide valuable biological information for prioritizing particular sites of functional importance (*e*.*g*., in regulatory regions).

**Table 1.**
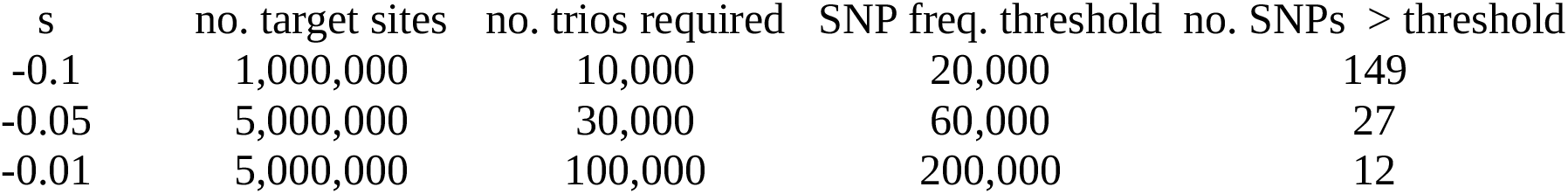
Segregation of strongly deleterious variants in simulations. For each selection coefficient, we show the number of target sites simulated, the number of trios required for accurate inference by TIDES and the number of SNPs segregating at frequency q ∼0.5 (shown as SNP count among parents).

| s | no. target sites | no. trios required | SNP freq. threshold | no. SNPs > threshold |
| --- | --- | --- | --- | --- |
| -0.1 | 1,000,000 | 10,000 | 20,000 | 149 |
| -0.05 | 5,000,000 | 30,000 | 60,000 | 27 |
| -0.01 | 5,000,000 | 100,000 | 200,000 | 12 |

#### Inferring the selection coefficient of single beneficial SNPs

While strongly deleterious SNPs tend to segregate in very small numbers even in large samples, the opposite is true for beneficial variants, which are subject to positive selection. Although the fraction of the genome where mutations are expected to be beneficial is considerably smaller than for deleterious mutations, there are examples of SNPs putatively under strong positive selection in recent human history (Stern, Wilton, and Nielsen 2019; Mathieson 2020). To investigated whether TIDES could be used to infer the strength of positive selection acting on single SNPs, we simulated datasets where the frequency *q* of the beneficial allele among parents is ∼0.5, and we varied the number of informative trios (10,000, 30,000 or 100,000), as well as *s* (0.01, 0.05 or 0.1) and *h* (0.5 or 1.0). We opted for including fully dominant beneficial alleles in order to explore TIDES’s power in this typically less explored (but still plausible) scenario. When performing inference, we flipped the sign of the prior on *s* (log-uniform in the open interval (10^−4^ ; 10^0^)). As expected, our statistical power is very similar to the analogous analysis of negative selection. TIDES is accurate in all scenarios of *s* = 0.1, whereas it requires 30,000 informative trios to have high accuracy when *s* = 0.05 and does not start to perform well until 100,000 trios and dominant selection for *s* = 0.01 (**Figure 5**). Since such variants are expected to segregate at a range of frequencies on their way to fixation (depending on their age and fitness effect), finding the necessary number of informative trios should not require that entire populations are sequenced, expediting the application of TIDES to study candidate beneficial SNPs. A particularly attractive application of our method will be to test whether variants with a strong signal of positive selection in the last few thousand years of human evolution are still under strong selection today. Lactase persistence is a canonical example of such recent and strong positive selection, (Bersaglieri et al. 2004), with the 13910*T variant segregating at frequencies of ∼0.77 and ∼0.43 in Northern and Southern Europeans, respectively (Liebert et al. 2017). Therefore, a total of ∼40,000 random trios from Northern Europe or ∼25,000 random trios from Southern Europe would be required to find ∼10,000 informative trios in each of these populations. Assuming that the children are old enough such that the advantage conferred by the derived allele has had the opportunity to be manifested, with these numbers, it will already be possible to test the hypothesis that the strength of (population-specific) ongoing selection is ∼0.1 or greater.

**Figure 5:**
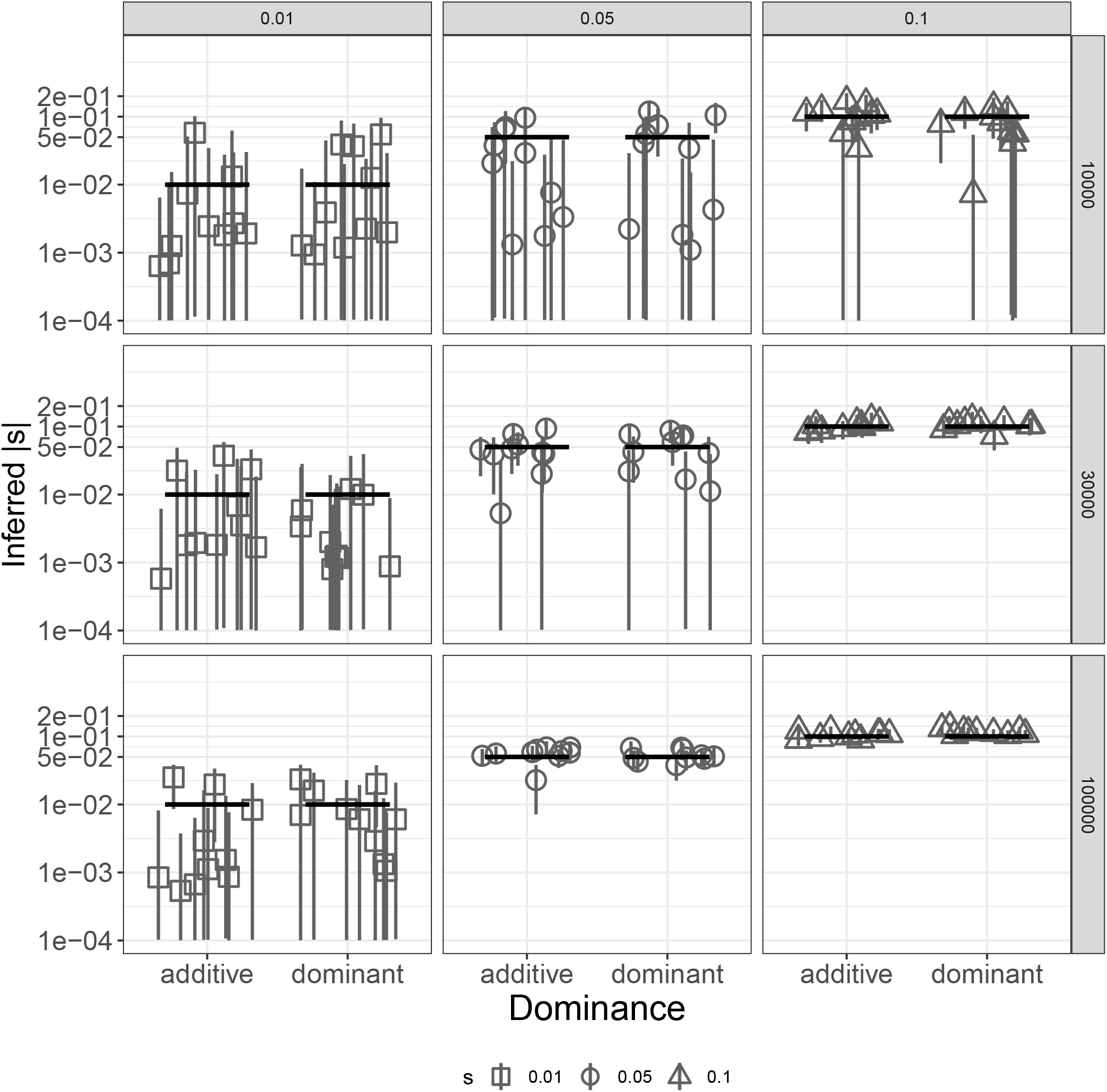
Inference of *s* from a single beneficial SNP with different dominance effects. Each scenario includes the estimates from 10 simulated datasets. Columns show different values of the true selection coefficient, and rows show different sample sizes, in terms of the number of trios used. True values are shown as black horizontal segments, with medians of the inferred posterior distributions denoted by gray shapes and their 95% credible intervals by gray vertical lines. Y-axis in log_10_ scale.

#### Inferring the dominance coefficient

In its pursuit of estimating selection coefficients, population genetics has paid less attention to the dominance effect of mutations, often assuming additivity (*h* = 0.5) (Eyre-Walker, Woolfit, and Phelps 2006; Boyko et al. 2008; Galtier 2016; Kim, Huber, and Lohmueller 2017; Tataru et al. 2017). But how mutant and wild-type alleles interact to influence fitness in heterozygous genotypes is crucial for understanding the evolutionary fate of alleles as well as their impact on genetic load and individual health. While *s* and *h* are mutually unidentifiable with SFS data (Huber et al. 2018), modeling the transmission of genotype counts in family trios allows us to tease them apart because *h* directly impacts the expected number of heterozygous but not homozygous sites in the children (**equation 1**). When inferring posterior distributions for the dominance coefficient using the exome-wide data from the simulations described above, we observe similar trends in accuracy as for the selection coefficient: the posterior distributions fall near the true values of *h* in both the strong and moderate selection scenarios, but not for weak selection due to the small value of |*n* * *s*| since the strength of selection also affects our ability to infer *h* from SNP transmission distortions (**Figure 6**). On the other hand, estimating the dominance coefficient for single variants is more challenging than estimating the selection coefficient in terms of the required sample size. For both negative selection (**Figure 7**) and positive selection (**Figure 8**), posterior distributions of *h* are too wide when |s| = 0.01 (spanning almost the entire range of the prior) whereas we need >30,000 informative trios when |s| = 0.1 and >100,000 informative trios when |s| = 0.05 for accurate inference.

**Figure 6:**
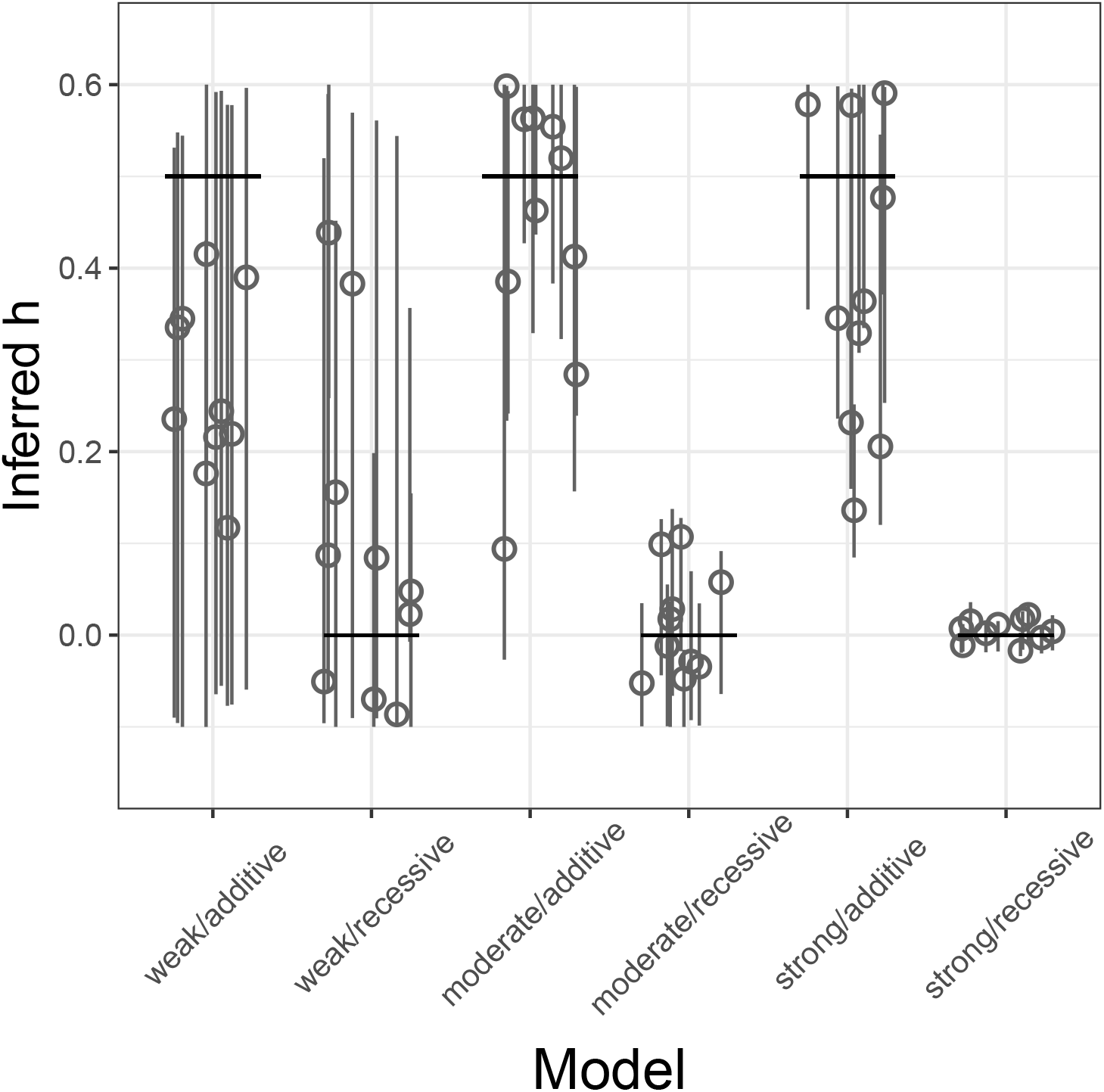
Inference of *h* from a genome-wide set of deleterious SNPs for different strengths of selection and dominance effects. Each scenario includes the estimates from 10 simulated datasets. True values are shown as black horizontal segments, with medians of the inferred posterior distributions denoted by gray circles and their 95% credible intervals by gray vertical lines. Each scenario includes 50,000 trios.

**Figure 7:**
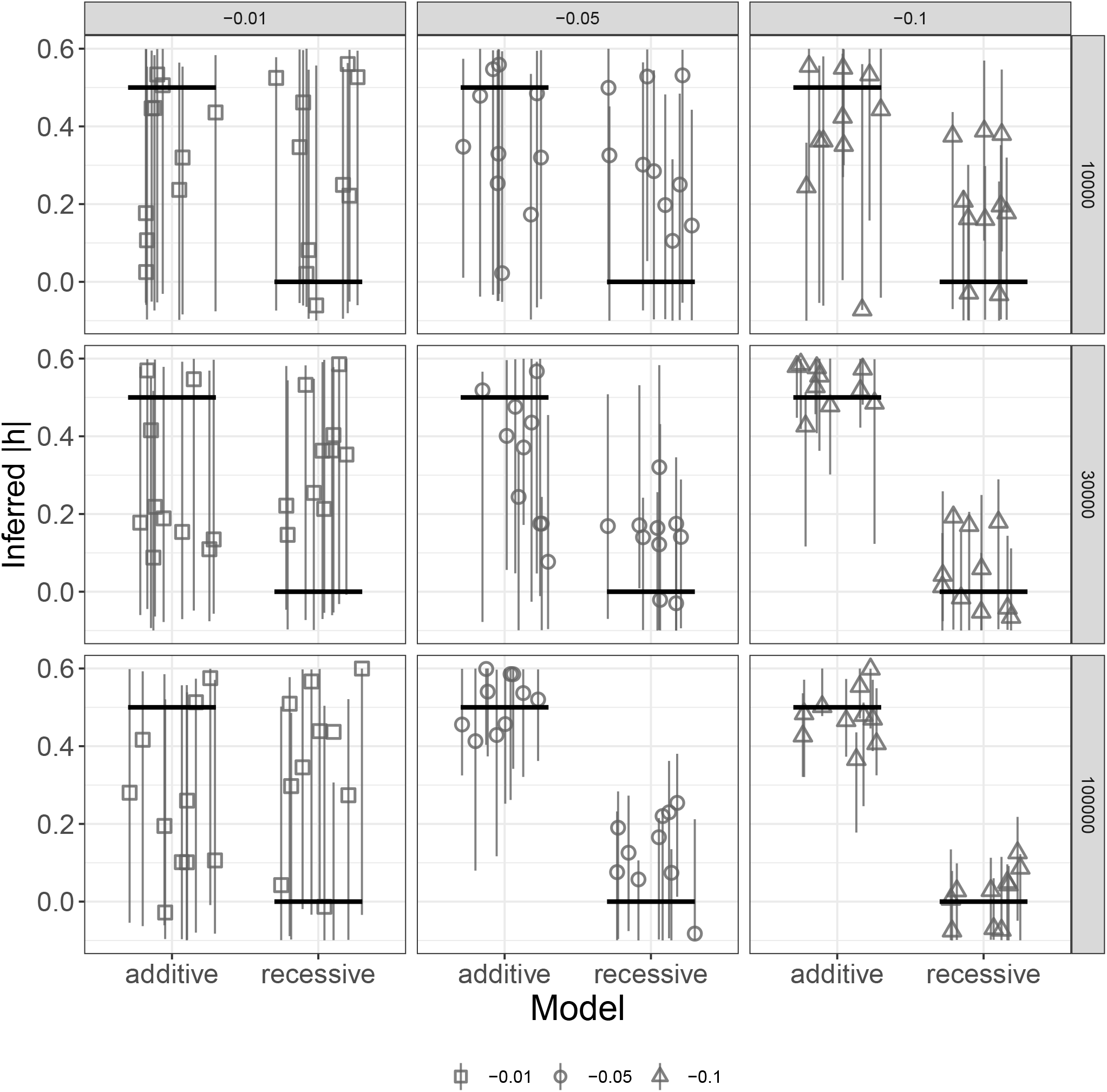
Inference of *h* from a single deleterious SNP with different dominance effects. Each scenario includes the estimates from 10 simulated datasets. Columns show different values of the true selection coefficient, and rows show different sample sizes, in terms of the number of trios used. True values are shown as black horizontal segments, with medians of the inferred posterior distributions denoted by gray shapes and their 95% credible intervals by gray vertical lines.

**Figure 8:**
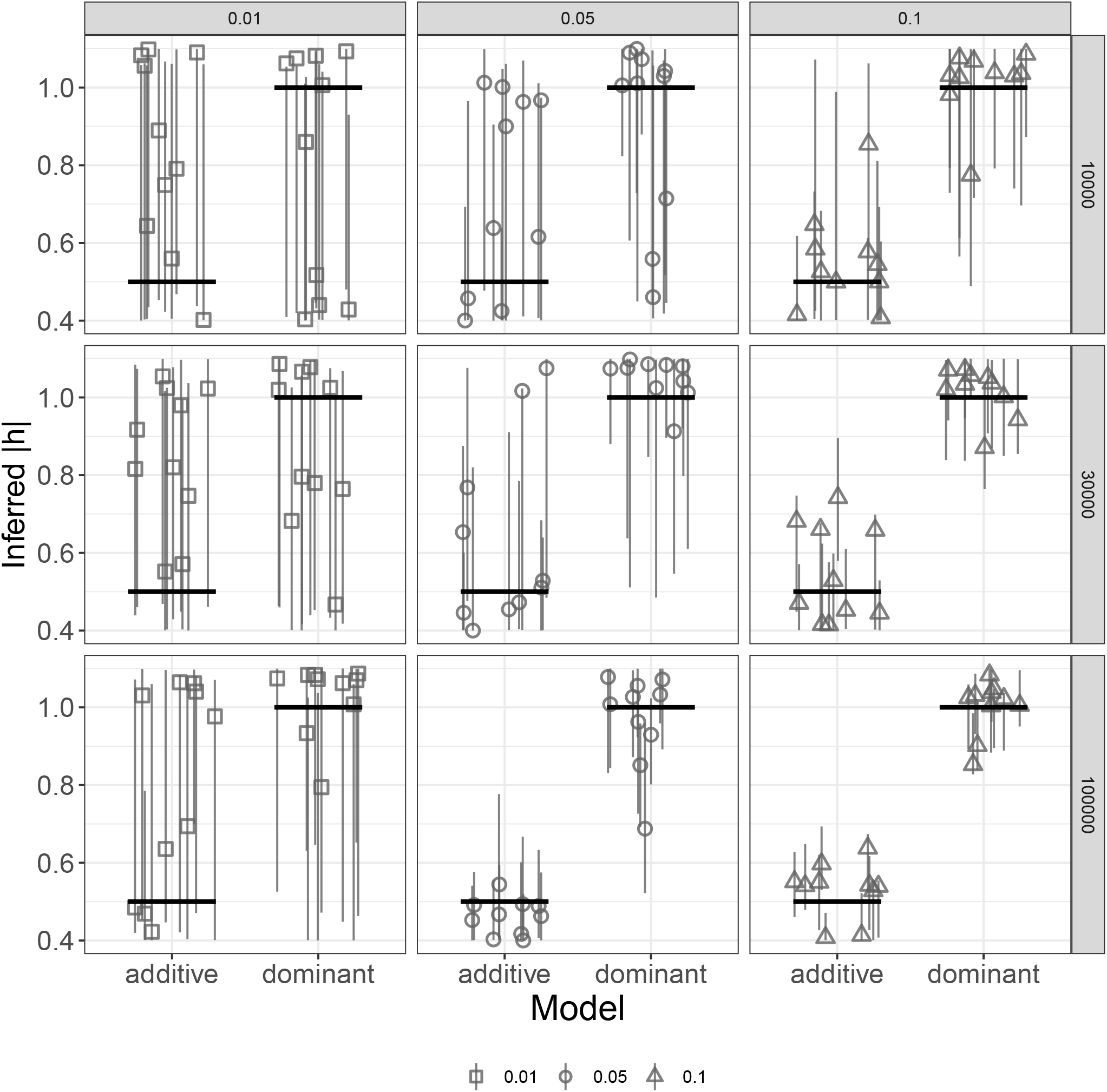
Inference of *h* from a single beneficial SNP with different dominance effects. Each scenario includes the estimates from 10 simulated datasets. Columns show different values of the true selection coefficient, and rows show different sample sizes, in terms of the number of trios used. True values are shown as black horizontal segments, with medians of the inferred posterior distributions denoted by gray shapes and their 95% credible intervals by gray vertical lines.

In addition to inferring posterior distributions for *h*, we can compare the fit of different models of dominance through Bayesian model selection (Csilléry et al. 2010; Csilléry, François, and Blum 2012). To this end, we fitted constrained models to each exome-wide dataset, then computed posterior probabilities for each model based on their acceptance rates in the rejection algorithm. We tested TIDES’ ability to distinguish between ‘neutral’, ‘additive’ and ‘recessive’ models, all of which carry historical meaning in population genetics. In the ‘additive’ and ‘recessive’ models *h* is fixed to 0.5 and 0, respectively, and only *s* is drawn from its prior distribution. In the ‘neutral’ model, *s* is fixed to 0 and any fluctuation in the frequency of SNPs is due to genetic drift alone. To benchmark the accuracy of our method in model selection, we ascribed equal prior probabilities to the three models (in ABC, we do this by considering the same number of candidate simulations under each model). Using this framework, TIDES shows remarkable accuracy in classifying data sets (**Figure 9**), with posterior probabilities of 1.0 being assigned to the correct model in all 20 replicates of the strong selection regime. Likewise, posterior probabilities of 1.0 are assigned to ‘recessive’ in all 10 replicates of recessive & moderate selection whereas posterior probabilities > 0.9 are assigned to ‘additive’ in all the 10 replicates of additive & moderate selection. As seen above, the weak selection regime is the most challenging, where the posterior probabilities are diffuse across the three models in all 20 replicates. Here, TIDES often cannot reject a neutral model of evolution, in agreement with sample size being insufficient for efficient negative selection. In summary, our model offers high accuracy and precision to distinguish between recessive and additive models of selection.

**Figure 9:**
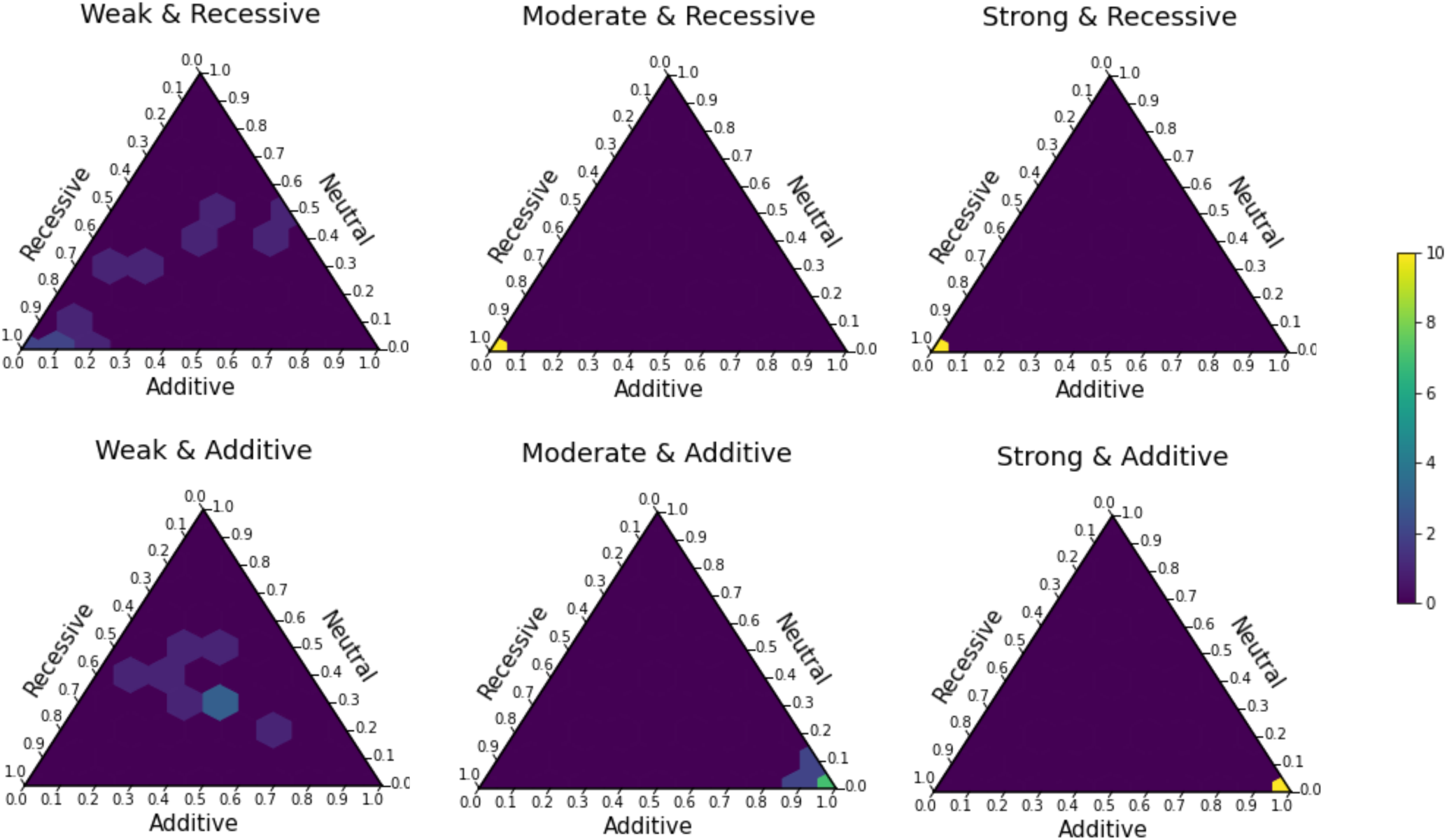
TIDES can accurately distinguish among neutral, recessive, additive models of selection. True models are shown above each simplex. Coordinates along each axis denote posterior probabilities assigned to the respective model. Color of each tile represents the proportion of simulations that fall within that probability bin (scale shown in far right).

### Implementation and software availability

We opted to embed TIDES in an ABC framework because its high flexibility will foster extensions of the model in the future. ABC has been employed to study a wide range of complex problems in population genetics, where an expression for the likelihood is bypassed by simulating the evolutionary process extensively (Beaumont 2019; Csilléry et al. 2010). The idea is to generate data according to parameter values drawn from prior distributions and retain those that best approximate the observed data, according to a suitable set of summary statistics that reduces the dimensionality of the data. In the Bayesian paradigm, the resulting posterior distributions represent our updated knowledge about the parameters of interest. Crucial to the performance of ABC methods is (1) the ability to simulate the data-generating process with high fidelity and speed (in order to build a large number of datasets that will be used to “train” the model); and (2) access to summary statistics that contain enough information about the parameters (in order to objectively score the similarity between simulated and observed data).

Up to a limit, the accuracy and precision of ABC methods depend on the total number of simulations performed (Beaumont 2019). To optimize our sampling scheme of parameter values, we implemented a two-step procedure (**Algorithm 1** and **Algorithm 2**): 1) TIDES executes a pilot set of simulations and employs the rejection algorithm to generate posterior distributions for *s* and *h*; 2) it uses samples from such posteriors as empirical priors for a second and larger round of simulations, which is then forwarded downstream to perform the final inference in R using regression adjustment techniques (Beaumont, Zhang, and Balding 2002; Csilléry, François, and Blum 2012; Blum 2018). This approach follows naturally from the Bayesian paradigm, adding information to the prior distributions while allowing full parallelization of the parameter draws within each step, and is similar in spirit to population Monte Carlo (Cappé et al. 2004; Sisson, Fan, and Tanaka 2007). As a result, TIDES’ speed increases almost linearly with the number of computing threads. Naturally, the choice of prior also reflects on the accuracy and precision of the posterior distributions. In our simulation study, we used uniform, weak priors to assess how much information our model can extract from the data, and in this sense our results were somewhat conservative with respect to accuracy. In principle, however, one can specify any arbitrary distribution as priors for *s* and *h*. A public link to the TIDES software package (composed of the simulation engine written in C++ and R scripts for downstream analyses and visualization of the results) will be available upon publication.

## DISCUSSION

Sequence data from family trios offers a new perspective for the inference of natural selection. We have implemented these ideas into a new statistical model called TIDES, which has several conceptual improvements over traditional SFS-based methods. First, the parent-offspring structure grants immunity to biases arising from a complex demographic history (Bates et al. 2020). Demography *sensu lato* (including population size changes, sub-division and migration) has been shown to be a strong confounder in the inference of selection in general (Nielsen et al. 2007; 2009; Williamson et al. 2005), and eliminating this effect consists a major enhancement in model design. Second, TIDES does not require the specification of putatively neutral variants as a measure of contrast. In SFS-based methods these are usually represented by synonymous SNPs, which is sub-optimal because synonymous SNPs may be themselves under selection for codon usage (Plotkin and Kudla 2011). Third, we explicitly model linkage and selective interference among SNPs, which is particularly relevant in regions of low recombination rate. Fourth, TIDES jointly infers the dominance coefficient *h*. This not only improves inference of *s* by integrating it over a range of dominance values, it also enables directly testing for additive *vs* recessive effects of mutations, a notoriously challenging problem in human population genetics because *h* and *s* are unidentifiable in the SFS (Huber et al. 2018). Methods to infer strong negative selection must rely on the assumption of mutation-selection balance (Haldane 1937; Weghorn et al. 2019), hence can only infer the strength of selection against the heterozygote genotype (Cassa et al. 2017). Finally, TIDES is notably sensitive to the strength of selection acting on the current generation. This is an unique feature of our method that avoids conflating the selective constraint from different periods in time into a single estimate. For instance, it is conceivable that human cultural evolution (the advent of medicine in particular) may have modulated the selective pressure on several genes, and our method captures such updated state of selective constraint. These improvements come at the cost of targeting inference at the average *s* rather than a DFE (when using multiple SNPs) and requiring sample sizes of the order of tens of thousands of trios. Fortunately, extravagantly large genomic datasets are becoming commonplace, and we anticipate that TIDES will make its debut analyzing human data within the next few years. Application to domesticated species will follow, where breeding and genetic engineering are employed to impose changes along phenotypic gradients.

There are yet other properties that distinguish TIDES from existing methods. First, because TIDES computes summary statistics conditioning on the parental haplotypes, it is able to perform inference in biased data sets (*e*.*g*., where genetic load is higher than average), whereas existing methods require random samples from the population. Second, the type of polymorphism analyzed is not restricted to SNPs. It is straightforward to infer the *s* and *h* from structural variants such as copy number variation and chromosomal inversions, some of which have already been hypothesized to have strong phenotypic and fitness effects (Alonge et al. 2020; Hämälä et al. 2021). Considering the flexibility that results from TIDES’ properties put together, we anticipate that within the next few years, it will fundamentally change inference of natural selection.

A thought-provoking possibility is that TIDES may open an avenue for experimental evolution in multi-cellular organisms with relatively long generation times. Our results suggest that it is possible to infer *s* and *h* of single SNPs artificially introduced in model organisms (*e*.*g*., by CRISPR (Ran et al. 2013)). By measuring fitness in a large sample but in a single generation (as opposed to small samples collected over hundreds of generations, as traditionally done in single-cell organisms), TIDES can be used to study fitness effects of mutations (e.g., (Sarkisyan et al. 2016)) in highly constrained genes where natural variation is too low for traditional methods to work. Much like sequencing itself, the ease and cost of genetically-editing model organisms is expected to greatly improve in the upcoming years, broadening the range of application of single-generation inference of selection.

Future progress in genomics depends on extracting information from large data sets where assumptions about the relatedness of individuals (or lack thereof) break down. We have demonstrated that going beyond random samples from a population allows statistical methods to capture signals that are both more subtle (*e*.*g*., with respect to time-scale) and more robust (requiring fewer assumptions about the data-generating process) than the current state-of-the-art. Therefore, we believe that upcoming studies should prioritize the inclusion of family trios in their sequencing efforts (Meyer et al. 2012), since they provide an overall richer data structure that can be exploited to infer present-day recombination rates (Halldorsson et al. 2019), mutation rates (Francioli et al. 2015) and now dominance as well as selection. We hope that as the TIDES framework continues to develop, it will also inspire other groups to consider how to leverage the future abundance of family trio data to infer other types of selection.

## METHODS

TIDES is a fast simulator of meiosis followed by selection. Here we outline a typical execution with default options. For each of *n* trios, TIDES simulates an array of 150 zygotes. The number of zygotes each parent generates is proportional to the number of children they have in the dataset, avoiding bias in the presence of siblings. Since meiosis is independent of *s* and *h*, the *n* arrays are pre-constructed and retained throughout the execution of the program (**Figure 1A**), substantially improving computational performance. TIDES then iteratively 1) draws *s* and *h* values from their prior distributions; 2) computes the fitness of all zygotes and samples one per trio with probability proportional to their fitness (**Figure 1B**); 3) computes summary statistics for the batch of “selected” zygotes (**Figure 1C, Algorithm 1**; note that sampling one zygote with probability proportional to its fitness is more efficient than sampling one zygote at random, imposing viability selection, and repeating this process until a zygote survives in each trio). Contrasting summary statistics computed from the simulated survivors (**Q**_**sim**_) with those computed from the actual children (**Q**_**obs**_) offers an objective approach to inference: values of *s* and *h* that generate substantial differences between **Q**_**sim**_ and **Q**_**obs**_ are discarded, while those that best agree are used to paint posterior distributions for the parameters. In the pilot run, we use an uniform prior (e.g., (− 0.1, 0.6)) for *h* and a log-uniform (e.g., (−10^−5^, -10^−1^)) prior for *s*, the latter in order to more frequently sample from regions of low selection coefficients that would otherwise not be sufficiently explored by an uniform prior.

### ABC implementation

Given sufficient summary statistics, the accuracy of ABC converges to that of full likelihood methods as the acceptance rate of proposed parameters decreases towards zero (Beaumont 2019). In practice, the performance of specific ABC methods is limited by the total number of simulations performed by their engine (Csilléry et al. 2010). Therefore, motivated by improving computational efficiency, several approaches have been developed to sample parameter values from a region of high posterior density (Fan and Sisson 2018). These mostly rely on proposing a new set of values conditional on the acceptance of previous proposals. Consequently, these approaches complicate parallelization of the simulation engine, being of most value when individual simulations are computationally expensive. When focusing on the one-generation interval between parents and offspring, however, each individual simulation is computationally cheap such that in TIDES we prioritized multi-threading over elaborate sampling techniques. After obtaining a reference table of the simulation using TIDES, all downstream analyses regarding model selection and parameter inference were performed using a combination of packages *abc, rethinking, scales, cowplot* and *tidyverse* in R 3.6 (scripts available in the GitHub repository).

The algorithms below describe the ABC implementation of TIDES.

#### Algorithm 1

1. *Setting up* Compute summary statistics **Q**_**obs**_ for the actual children; For each trio 1..*n*, generate 150 zygotes based on parental haplotypes and sex-specific recombination maps;
2. *Pilot simulations* For each pilot simulation 1..*M*_*PILOT*_:
  a. draw *s* and *h* from their prior distributions;
  b. for each trio 1..*n*, compute the fitness of zygotes 1..150;
  c. for each trio 1..*n*, sample one zygote with probability proportional to its fitness;
  d. compute summary statistics **Q**_**sim**_ for the sample on *n* selected zygotes;
3. *Updating priors* Accept a proportion *t* of the pilot simulations using the standard rejection algorithm based on the Euclidean distances between **Q**_**obs**_ and each **Q**_**sim**_; Set the sorted arrays of accepted *s* and *h* values as prior distributions for step 4;
4. *Final simulations* For each simulation 1..*M*_*FINAL*_:
  a. draw *s* and *h* from their updated prior distributions using **Algorithm 2**;
  b. for each trio 1..*n*, compute the fitness of zygotes 1..150;
  c. for each trio 1..*n*, sample one zygote with probability proportional to its fitness;
  d. compute summary statistics **Q**_**sim**_ on the sample on *n* selected zygotes;
5. Use TIDES output files as input for *abc_adjust*.*R* to paint the posterior distributions of *s* and *h* using rejection followed by regression adjustment.

#### Algorithm 2

1. Let **w**_**1**_ be an element drawn uniformly at random from the sorted array of parameter values;
2. If **w**_**1**_ is the first element of the array, let **w**_**2**_ be the next element; Else if **w**_**1**_ is the last element of the array, let **w**_**2**_ be the previous element; Else set **w**_**1**_ and **w**_**2**_ as the previous and next elements in the list, respectively;
3. Draw a random number uniformly between **w**_**1**_ and **w**_**2**_.

### Simulation study

When benchmarking inference on individual SNPs, genomes for the test datasets were simulated trivially within TIDES itself – each of the two haplotypes in each parent received the derived allele with 50% probability. Then, when joining females and males in couples, we avoided matches where both parents were homozygous for the same allele. When benchmarking inference on a large set of candidate SNPs (*e*.*g*,, all non-synonymous mutations), the simulation of the test dataset was more involved. Parental genomes were generated using SLiM 3.1 (Haller and Messer 2018). The sequences were 66.8 Mb in size, approximately the size of the human exonic coordinates obtained with Ensembl annotation in the *biomaRt* package. Non-synonymous sites were distributed according to exome coordinates provided for GRCh38 in Ensembl, after removing overlapping genes. The non-synonymous mutation rate was set to 6.65e-09 per site per generation and we used sex-specific recombination maps from deCODE (Halldorsson et al. 2019). The non-synonymous mutation rate is in the low end of the spectrum commonly adopted, meaning that our simulations are conservative with respect to the amount of standing deleterious diversity and therefore our simulation study is likewise conservative with respect to statistical power. The demography of the sample approximated the demography of Europeans (Gravel et al. 2011), where we omitted African populations for computational efficiency as well as increased the number of generations from 58,000 to 58,300 so that ∼60,000 diploid individuals are sampled in present time. To allow reproducibility, we set the random seed of simulations in each evolutionary scenario to its corresponding replicate number (1-10). All scripts necessary to reproduce the above procedures will be found in the GitHub repository upon publication.

To follow the human demographic model precisely and generate a realistic and predictable amount of standing genetic variation, the simulations above were carried out under the Wright-Fisher model that uses relative fitness among individuals and where selection occurs in the mating stage of the life cycle. Because we focused on viability selection, we had SLiM output genomes from the parental generation exclusively. These were then input in TIDES where they underwent viability selection followed by random pairing of females and males, reproduction, and finally viability selection was imposed on the resulting embryos. These trios became the “observed” data in each of our simulated scenarios. To ensure that in all scenarios exactly 50,000 children are available after viability selection in the embryos (*i*.*e*., so we can directly relate statistical power to sample size), we over-shot and generated 10 embryos per couple, subsequently down-sampling the number of surviving children to 50,000 at random. Since for some couples more than one child survives this process, our simulated data sets naturally contain siblings. Viability selection was executed according to the description in the SLiM manual (individuals where deleted if their fitness was smaller than a uniform random number between 0 and 1) and using the same *s* and *h* values as in the Wright-Fisher step of the simulation, except for simulated data sets where the purpose was to test sensitivity to recent changes in *s* (**Figure 3b**). For the data sets where the selection coefficient of each mutation follows a DFE, this information was extracted from the VCF files output by SLiM and the fitness of each individual was computed using its exact genotype-fitness map. In summary, the simulated trios experienced a Wright-Fisher human demographic model with mating selection, followed by non-Wright-Fisher dynamics with viability selection in the last two generations.

For the simulations to assess the frequency of strongly deleterious variants segregating in the population (**Table 1**), we used a simpler demographic model where the population size is constant at 10,000 individuals for 57,000 generations and then grows for 1,000 generations until it reaches 10,000,000. In these simulations, the number of target sites is 1,000,000 (for s = -0.1); 5,000,000 (for s = -0.05); and 5,000,000 (for s = -0.01), with a constant recombination rate of 10^−3^ between each pair of sites in order to partially mimic their dispersion across the genome. Simulations to benchmark single-SNP inference were conducted within TIDES itself by assigning the derived allele to each parental haplotype according to a Bernoulli trial with p = 0.5. When performing inference on (single) beneficial SNPs, we subtracted 0.2 from the fitness of each individual throughout the ABC simulations to prevent individuals from having survival probabilities > 1.

In our simulation study of human-like exons, we omitted *de novo* mutations (in the last generation) from our test data in order to focus on the reduction of deleterious variation caused by negative selection between parents and offspring. However, we provide two options for TIDES users to accommodate *de novo* mutations in real data sets. First, one can specify the total number of sites that can be targeted by deleterious mutations (L) as well as the mutation rate per site per generation among these sites (μ). In this case, SNPs are added to each simulated zygote with Poisson rate equal to *L*×*μ*. In case the user-specified rate is zero (its default value), *de novo* mutations are identified as those absent from parents but present in children, from which they are removed. These options are presented in the test run that can be found in TIDES GitHub page, which will become public upon acceptance of the manuscript.

---

**Figure S1:**
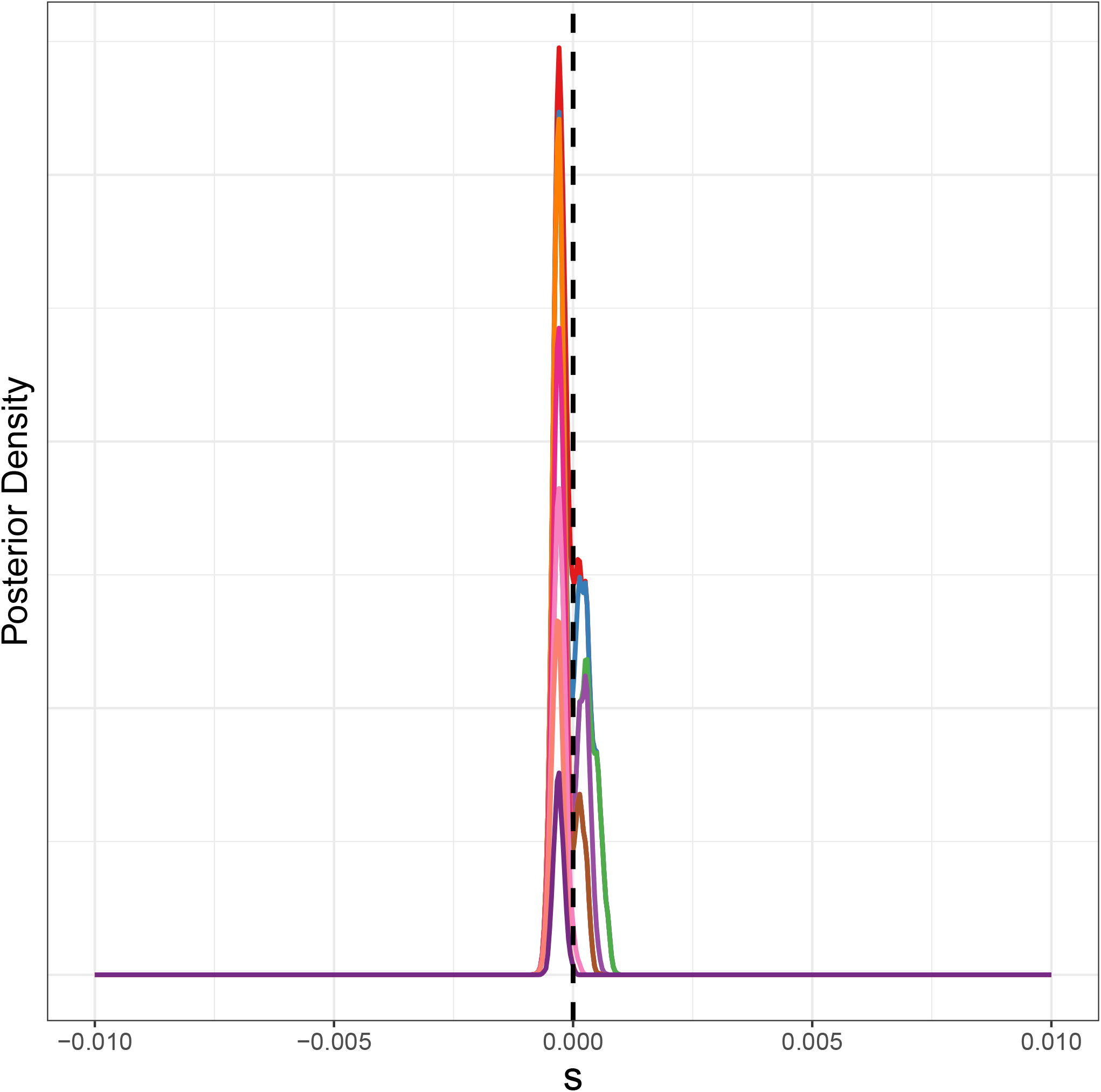
Posterior distributions of *s* under neutrality. Each color represents the posterior density inferred from one of 10 replicate scenarios. Here 50,000 trios are used.

## ACKNOWLEDGMENTS

We are thankful to Armaan Singh, Chris Kyriazis, Fabian Klötzl, Flora Jay, Jesse Garcia, Julien Dutheil and Nandita Garud for discussions about this work. This work was supported by the National Institutes of Health grant R35GM119856 to KEL.

